# Identification of simplified microbial communities that inhibit *Clostridioides difficile* infection through dilution/extinction

**DOI:** 10.1101/2020.04.23.058867

**Authors:** Jennifer M. Auchtung, Eva C. Preisner, James Collins, Armando I. Lerma, Robert A. Britton

**Affiliations:** Alkek Center for Metagenomics and Microbiome Research and Department of Molecular Virology and Microbiology, Baylor College of Medicine, Houston, TX USA; Nebraska Foods for Health Center and Department of Food Science and Technology, University of Nebraska-Lincoln, Lincoln, NE USA; Department of Microbiology & Immunology, University of Louisville School of Medicine, Louisville, KY USA; Department of Food Science and Technology, University of Nebraska-Lincoln, Lincoln, NE USA

**Author notes:** **Corresponding author:** Robert A. Britton.

## Abstract

The gastrointestinal microbiome plays an important role in limiting susceptibility to infection with *Clostridioides difficile*. To better understand the ecology of bacteria important for *C. difficile* colonization resistance, we developed an experimental platform to simplify complex communities of fecal bacteria through dilution and rapidly screen for their ability to inhibit *C. difficile in vitro*. We simplified complex communities from six fecal donors and found that 17% of simplified communities inhibited *C. difficile* growth when initially isolated and when re-cultured from frozen stocks. Composition varied between simplified communities based upon fecal donor used for dilution; complexity ranged from 19-67 OTUs. One simplified community could be further simplified through dilution and retain the ability to inhibit *C. difficile*. We tested efficacy of seven simplified communities in a humanized microbiota mouse model and found that four communities were able to significantly reduce the severity of the initial *C. difficile* infection and limit susceptibility to disease relapse. Analysis of fecal microbiomes from treated mice demonstrated that simplified communities accelerated recovery of endogenous bacteria and led to stable engraftment of at least 20% of bacteria from simplified communities. Overall, the insights gained through the identification and characterization of these simplified communities increase our understanding of the microbial dynamics of *C. difficile* infection and recovery.

**Importance:** *Clostridioides difficile* is the leading cause of antibiotic-associated diarrhea and a significant healthcare burden. While fecal microbiota transplantation is highly effective at treating recurrent *C. difficile* disease, uncertainties about the undefined composition of fecal material and potential long-term unintended health consequences have motivated studies to identify new communities of simple microbes that will be effective at treating disease. This work describes a platform for rapidly identifying and screening new simplified communities of microbes for efficacy in treating *C. difficile* infection and identifies four new simplified communities of microbes with potential for development of new therapies to treat *C. difficile* disease in humans. While this platform was developed and validated to model infection with *C. difficile*, the underlying principles described in the paper could be easily modified to develop therapeutics to treat other gastrointestinal diseases.

## Introduction

*Clostridioides difficile* is the most common cause of antibiotic-associated diarrhea (1-3). Estimates of annual healthcare costs associated with treating *C. difficile* infection (CDI) in the US range from $1 – 4.8 billion (4). Although uncomplicated infections are typically self-limiting, severe infections require treatment (5). For approximately 25% of patients, resolution of primary infection is followed by one or more rounds of recurrent *C. difficile* infection (rCDI; (6)), which diminishes quality of life and contributes to overall healthcare costs (7, 8).

To cause disease, ingested *C. difficile* spores must germinate into vegetative cells that produce toxins. The bile salt, cholate and its derivatives, stimulate *C. difficile* germination, along with co-germinants glycine and other amino acids (9, 10). The gastrointestinal microbiome plays key roles in limiting symptomatic CDI by competing with *C. difficile* for nutrients (11-13), producing metabolites that inhibit *C. difficile* growth (14-18), maintaining immune homeostasis (19-21) and metabolizing primary bile salts into secondary bile salts that inhibit the growth of vegetative cells (9, 22, 23). Antibiotic treatment leads to loss of GI microbiome diversity (24-26), is a key risk factor for primary infection (27-30), and contributes to susceptibility to recurrent disease.

Several different therapies are currently used to treat rCDI and act to limit different aspects of cell growth and pathogenicity. Clinical cure rates of 70% have been reported following extended-pulsed administration of the antibiotic fidaxomicin (31). An 80% cure rate has been reported in patients with primary and recurrent CDI treated with bezlotoxumab, an antibody that targets *C. difficile* Toxin B (32). Fecal microbiome transplantation (FMT) for treatment of rCDI has reported cure rates from 44-100% (33), (34). However, concern for potential adverse events (e.g., (35) motivates ongoing studies to develop alternatives to FMT to treat rCDI.

Defined community microbial therapeutics are one alternative to FMT under investigation. Previous studies have demonstrated success in administration of a consortium of 10 (36) or 33 (37) human fecal bacteria for treatment of rCDI. Despite these advances, no defined microbial therapeutic is currently available for treatment of CDI. One limitation to developing microbial community therapeutics is the availability of appropriate models for rapid screening. We developed a coupled *in vitro* and *in vivo* platform to screen simple microbial communities for their ability to prevent CDI. We identified four simplified communities that limited *C. difficile* growth *in vitro* and reduced severity of disease *in vivo*. While the potential of these communities to treat disease in humans is unknown, the approaches could be applied to identification of additional simplified communities to treat rCDI and microbiome-linked diseases.

## Results

### Identification of simplified communities that limit *C. difficile* growth *in vitro*

To identify simplified communities that could suppress *C. difficile*, we applied a dilution/extinction strategy (38, 39) as outlined in Fig. 1A. In this approach, complex fecal communities are simplified through dilution, with abundant organisms preserved and rare organisms randomly distributed or lost as predicted by the Poisson distribution. Complex communities were established in minibioreactor arrays (MBRAs) from six individual fecal donors and allowed to stabilize. Cell density was measured and used with published OTU abundance data (40) to estimate dilutions needed to simplify communities by 25-60%. Following dilution, simplified communities were stabilized in continuous culture before challenge with 10^4^ vegetative *C. difficile* cells. By measuring *C. difficile* proliferation in each bioreactor over time, we identified nine highly suppressive simplified communities that lowered *C. difficile* levels ≥10,000 times and 15 moderately suppressive communities that lowered *C. difficile* levels by ≥ 100 times lower compared to *C. difficile* cultivated alone in bioreactors (Fig. 1B).

**Fig 1.**
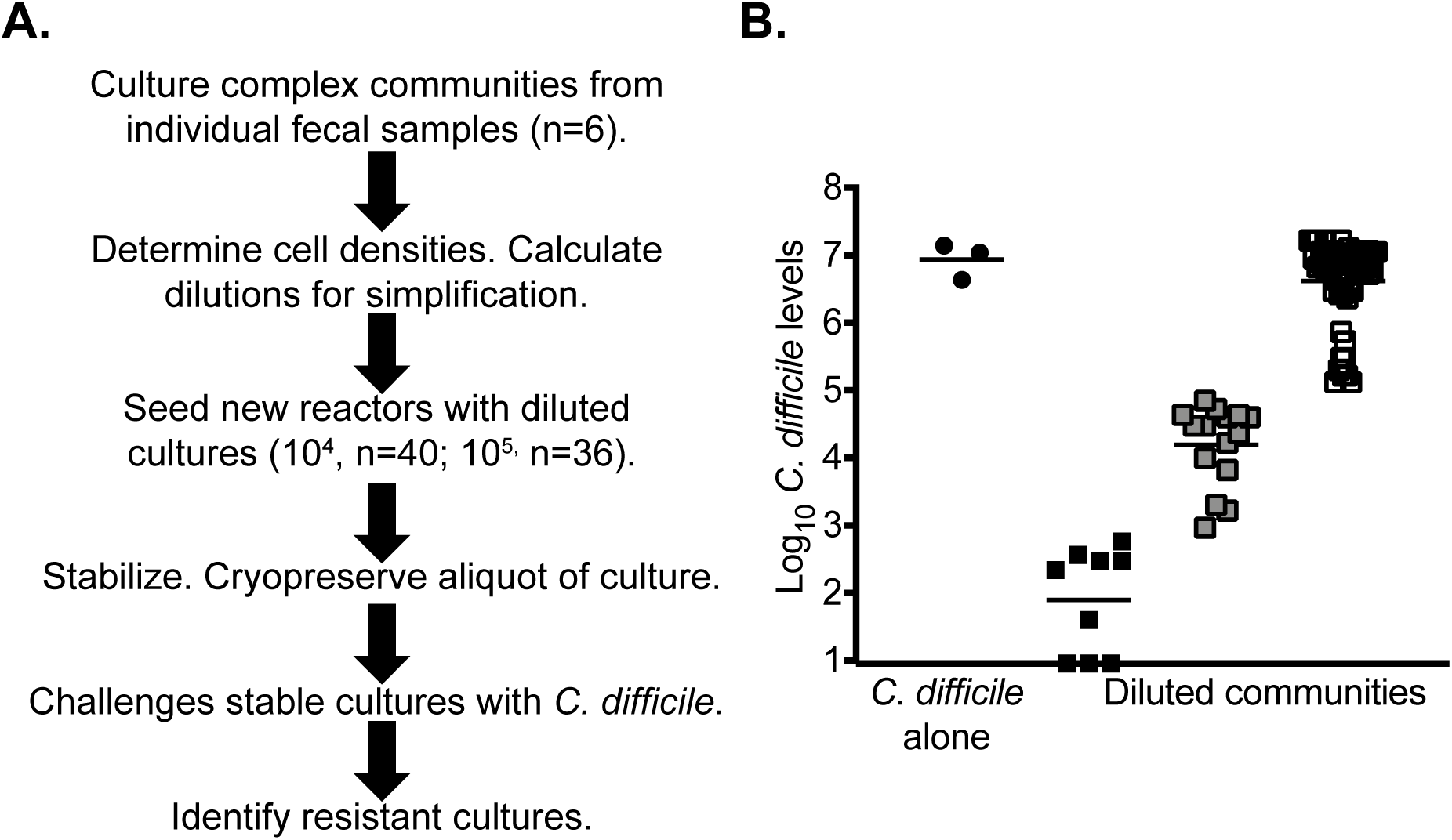
Identification of simplified communities that inhibit *C. difficile* proliferation through dilution/extinction community assembly. (A) Overview of process to identify simplified communities. (B) Log_10_ *C. difficile* levels measured in diluted communities on day 5/6 following challenge. Circles: cultures inoculated with *C. difficile* alone; Squares: stable diluted communities that suppress *C. difficile* by >10,000-fold (black squares), 100-10,000 -fold(gray squares), or by <100-fold (open squares) compared to growth of *C. difficile* alone. Lines represent the geometric means of the populations.

To better understand how dilution impacted community composition, we compared the 16S rRNA gene content between *C. difficile-*resistant complex and simplified communities. The median number of OTUs in the complex communities was 67; the median number of OTUs in 10^−4^ and 10^−5^ diluted communities were 50 and 42 (Fig. 2A). Microbial diversity was also reduced by dilution (Fig. 2B).

**Fig 2.**
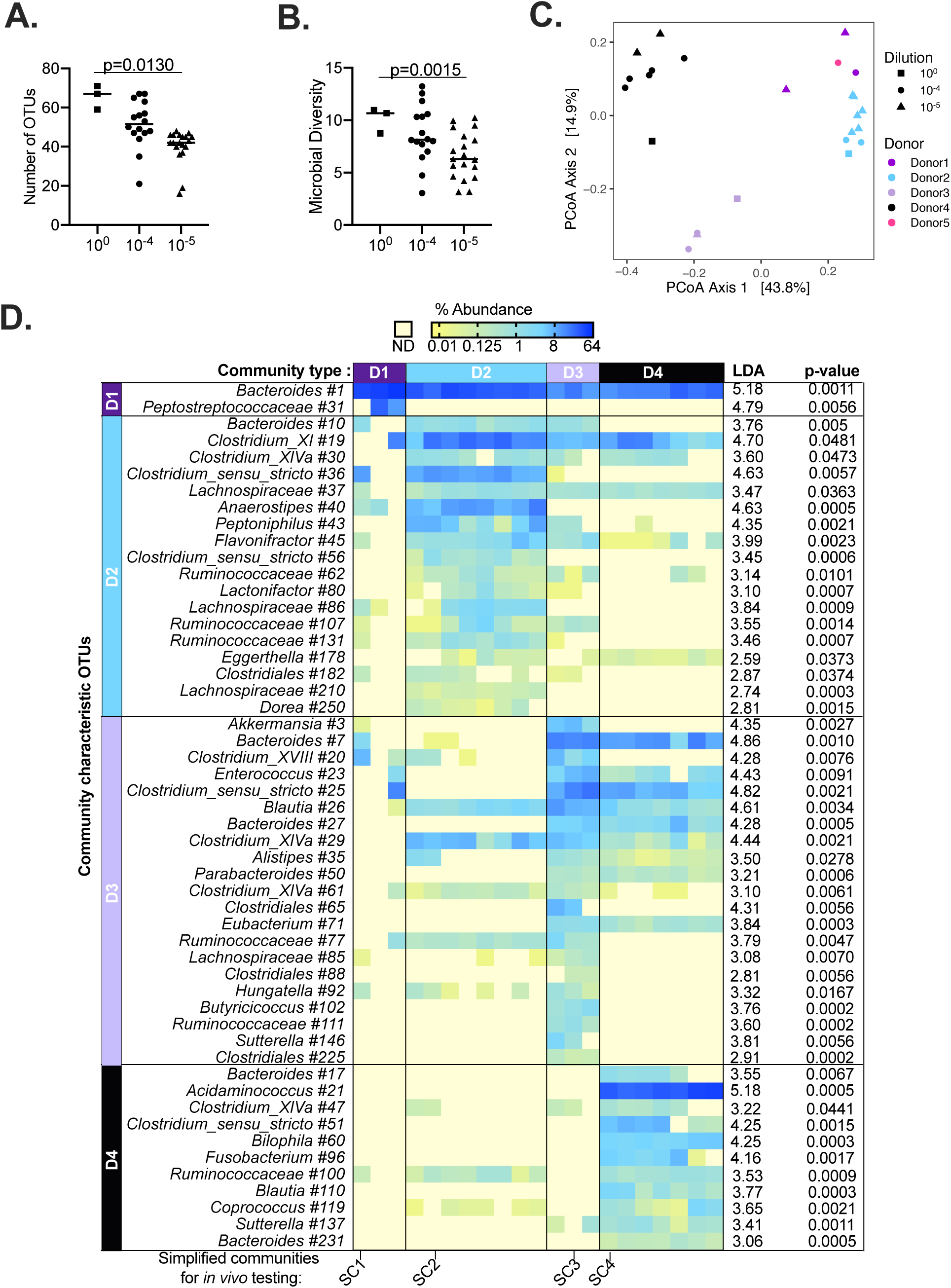
Comparison of microbial communities present in complex fecal donor and simplified communities that inhibit *C. difficile* growth. (A) The number of OTUs and (B) microbial diversity (Inverse Simpson) detected in complex communities from fecal donors (10^0^, squares), 10^−4^ (circles), and 10^−5^ (triangles) diluted communities was plotted. Lines represent medians; p-values <0.05 as calculated by one-way Kruskal-Wallis testing with Dunn’s correction for multiple comparisons are reported. (C) Principle Coordinates Analysis (PCoA) visualization of Bray-Curtis dissimilarities between complex communities from fecal donors (10^0^, squares), 10^−4^ (circles) and 10^−5^ (triangles) diluted community samples. Colors represent different fecal donors as indicated. Percent of variation described by each axis indicated in parentheses. Permutational ANOVA provided strong support for segregation of simplified communities by fecal donor (F-statistic=13.44; R^2^=0.73; p-value=0.001). (D) Significant OTUs that differentiate between D1, D2, D3, and D4 diluted communities are organized by community for which they are characteristic as indicated. OTUs are classified to the lowest taxonomic level that could be confidently assigned (≥80% confidence). The percent abundance of each OTU was plotted across all samples, which are arranged by donor community type as indicated at the top of the figure. Values ranged from 0.01% (yellow) to 64% (dark blue) of total sequences as indicated; pale yellow indicates no detected sequences (ND). LDA scores and p-values are indicated to the right of the heat map. The representative sequence for *Peptostreptococcaceae* #31 was 100% identical to *C. difficile* 16S rRNA. Diluted communities selected for *in vivo* testing are indicated in below the heat map (SC1 = D1, SC2=D2, SC3=D3, SC4=D4).

### CDI-resistant simplified communities separate into distinct community types

We compared differences in community structure across communities and found that communities separated primarily by fecal donor used for dilution (Fig. 2C). We identified OTUs characteristic of differences between simplified communities diluted from donors 1-4 (Fig. 2D). Different *Bacteroides* OTUs were enriched in each of the D1-D4 communities; *Anaerostipes, Clostridium_sensu_stricto, Clostridium XI*, and *Peptoniphilus* were enriched in D2 communities; *Akkermansia, Blautia, Clostridium XVIII*, and *Enterococcus* were enriched in D3 communities; and *Acidaminococcus, Bilophila* and *Fusobacterium* were enriched in D4 communities.

### Simplified communities retain their ability to suppress *C. difficile* when re-cultured

We re-cultured 17 simplified communities from frozen stocks in triplicate MBRAs and allowed them to stabilize prior to challenge with *C. difficile*. 13 of the 17 communities were able to inhibit *C. difficile* upon re-culturing. Five communities suppressed *C. difficile* growth by ≥10,000-fold across all three replicates, two suppressed *C. difficile* by ≥100-fold across all three replicates, and six communities suppressed *C. difficile* in at least one replicate (Fig. S1). We selected one simplified community (SC) from each community type (SC1-SC4; Fig. 2D) that could suppress *C. difficile* when re-cultured to test in a mouse model of disease.

### SC1 and SC2 suppress *C. difficile*-associated disease (CDAD)

We tested SC1-SC4 for their ability to suppress CDAD in a humanized microbiota mouse (^HMb^mouse) model of disease (41). Two positive controls were used to test for suppression of CDAD: FMT freshly prepared from ^HMb^mice and a cryopreserved aliquot of human FMT previously used successfully in a CDI fecal transplant program (Savidge, personal communication). Fig. 3A depicts the strategy for testing SC1-SC4 in the mouse model. SC1-SC4 were re-cultured in continuous flow bioreactors. Mice were treated with antibiotics to disrupt the microbiome, then gavaged with cells from simplified communities, ^HMb^mouse or human FMT on three consecutive days; control mice were treated with vehicle (Phosphate Buffered Saline (PBS)). Body mass was measured daily beginning with the first day of gavage to test for potential toxicity of communities. Because mice treated with SC4 exhibited ∼5% body mass loss from baseline prior to *C. difficile* challenge (Fig. 3B), SC4-treated mice were excluded from further analyses.

**Figure 3.**
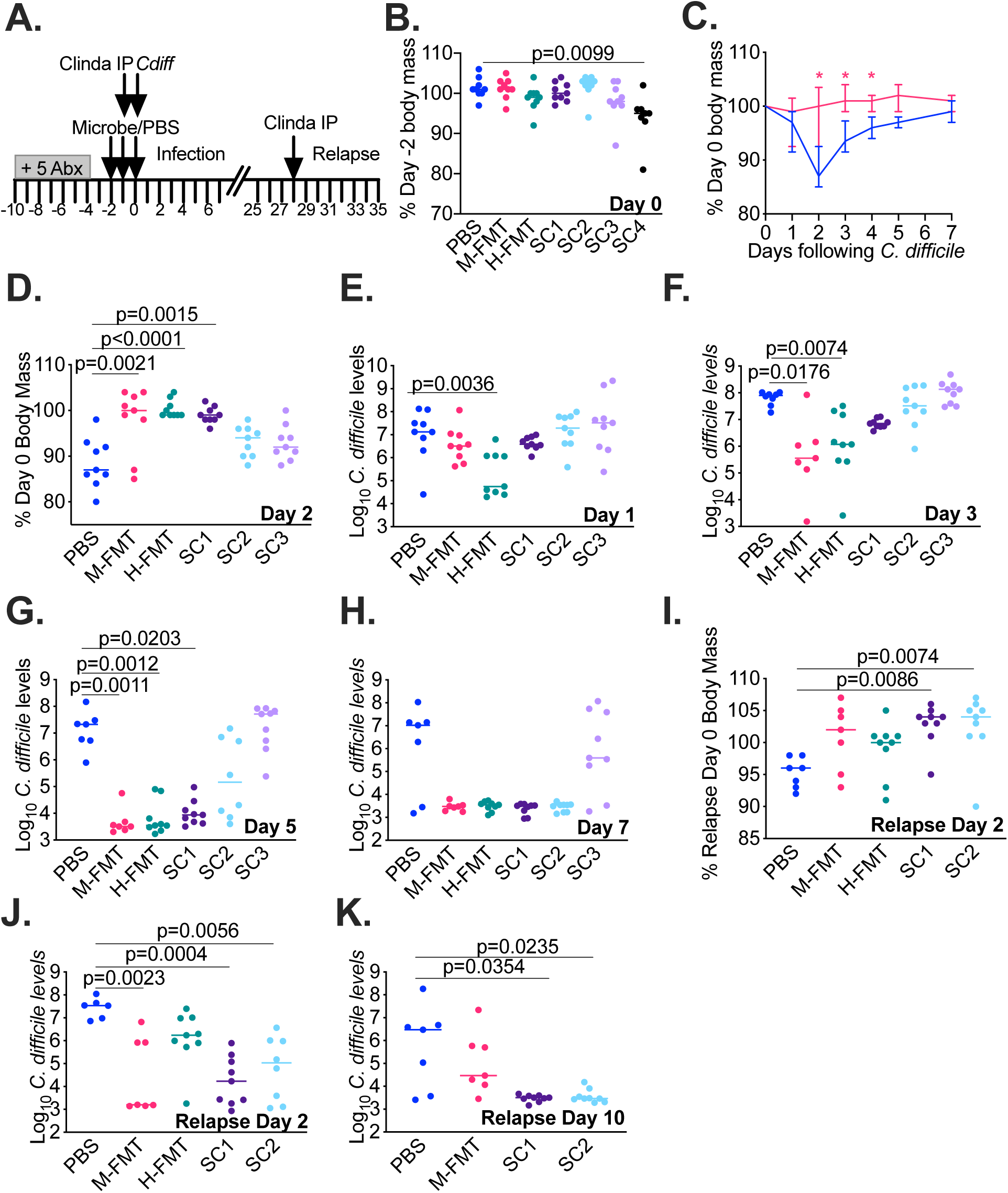
SC1 and SC2 suppress *C. difficile-*associated disease in ^HMb^mouse model. (A) Overview of infection and recurrence protocol used to evaluate simplified communities and FMT treatments. In (B)-(K), treatments indicated below the axis. Lines represent medians. Significance of differences between microbe and PBS-treated mice in each panel were evaluated with one-way Kruskal-Wallis testing with Dunn’s correction for multiple comparisons. p-values less than 0.05 are reported. (B) Percent of day -2 body mass on day 0 (prior to *C. difficile* challenge) following two days of treatment with simplified communities. (C) Percent of day 0 body mass of PBS and M-FMT treated mice over the first seven days of infection. Lines represent medians, error bars represent interquartile ranges, and asterisks indicate p-values <0.05. (D) Percent of day 0 body mass on day 2 following initial infection. Levels of *C. difficile* measured in the feces of treated mice on (E) day 1, (F) day 3, (G) day 5, or (H) day 7 following infection. (I) Percent of relapse day 0 body mass on relapse day 2. Level of *C. difficile* in mouse feces on (J) relapse day 2, (K) relapse day 7 and (L) relapse day 10. Two mice lost from PBS (days 3 and 4) and ^HMb^mouse FMT-treated groups (day 3) were included in calculations until death. Mice treated with SC3 were not tested for resistance to recurrent infection. *C. difficile* levels in H-FMT-treated mice were not tested on relapse day 10. Longitudinal data collected during initial infection and relapse are plotted in Fig. S2.

Following *C. difficile* challenge, mice treated with PBS exhibited up a decline in body mass (Fig. 3C-D) and shed *C. difficile* in their feces (Fig. 3E-H). In contrast, SC1-treated mice maintained their body mass, with levels similar to those observed in human or ^HMb^mouse FMT-treated mice (Fig. 3D) and exhibited more rapid clearance of *C. difficile* in feces (Fig. 3G-H). Trends towards reduced body mass loss in SC2 and SC3-treated and more rapid clearance of *C. difficile* in feces of SC2-treated mice (Fig. 3G-H) were not statistically significant (Fig. 3D).

Because *C. difficile* was cleared more rapidly in SC1 and SC2-treated mice, we tested whether these communities would reduce susceptibility to recurrent disease. Previously, we demonstrated relapse could be induced through a single IP injection of clindamycin (41). Four weeks following the initial *C. difficile* challenge, the majority of mice no longer shed *C. difficile* at detectable levels in their feces (Fig. S2N-Q). We treated mice with a single clindamycin IP injection, then measured changes in body mass and *C. difficile* levels. Consistent with relapse, we observed a modest body mass decline (Fig. 3I) and an increase of *C. difficile* in feces (Fig. 3J) of PBS-treated mice. In contrast, there was a modest body mass gain (Fig. 3K), reduced shedding of *C. difficile* in feces (Fig. 3J; Fig. S2P-Q) and more rapid clearance of *C. difficile* shed in feces (Fig. 3K; Fig. S2P-Q) in SC1 and SC2-treated mice following clindamycin IP. ^HMb^mouse and human FMT-treated mice had a more modest body mass gain that was not statistically significant from PBS-treated mice.

### SC2 can be further simplified and still inhibit *C. difficile* growth

We asked whether either SC1 or SC2 could be further simplified through dilution and retain the ability to prevent *C. difficile* infection. Cultures were diluted to a concentration of 250 CFU/ml (10^−6^ dilution); Poisson calculations indicated this dilution should reduce complexity of SC1 and SC2 to 17 and 31 OTUs, respectively. Newly diluted cultures were allowed to stabilize prior to challenge with *C. difficile*. Although 10^−6^ dilutions of SC1 lost the ability to inhibit *C. difficile*, 10^−6^ dilutions of SC2 continued to inhibit *C. difficile* growth (Fig. 4A). *C. difficile* inhibition was lost only when SC2 communities were diluted another 10-fold.

**Figure 4.**
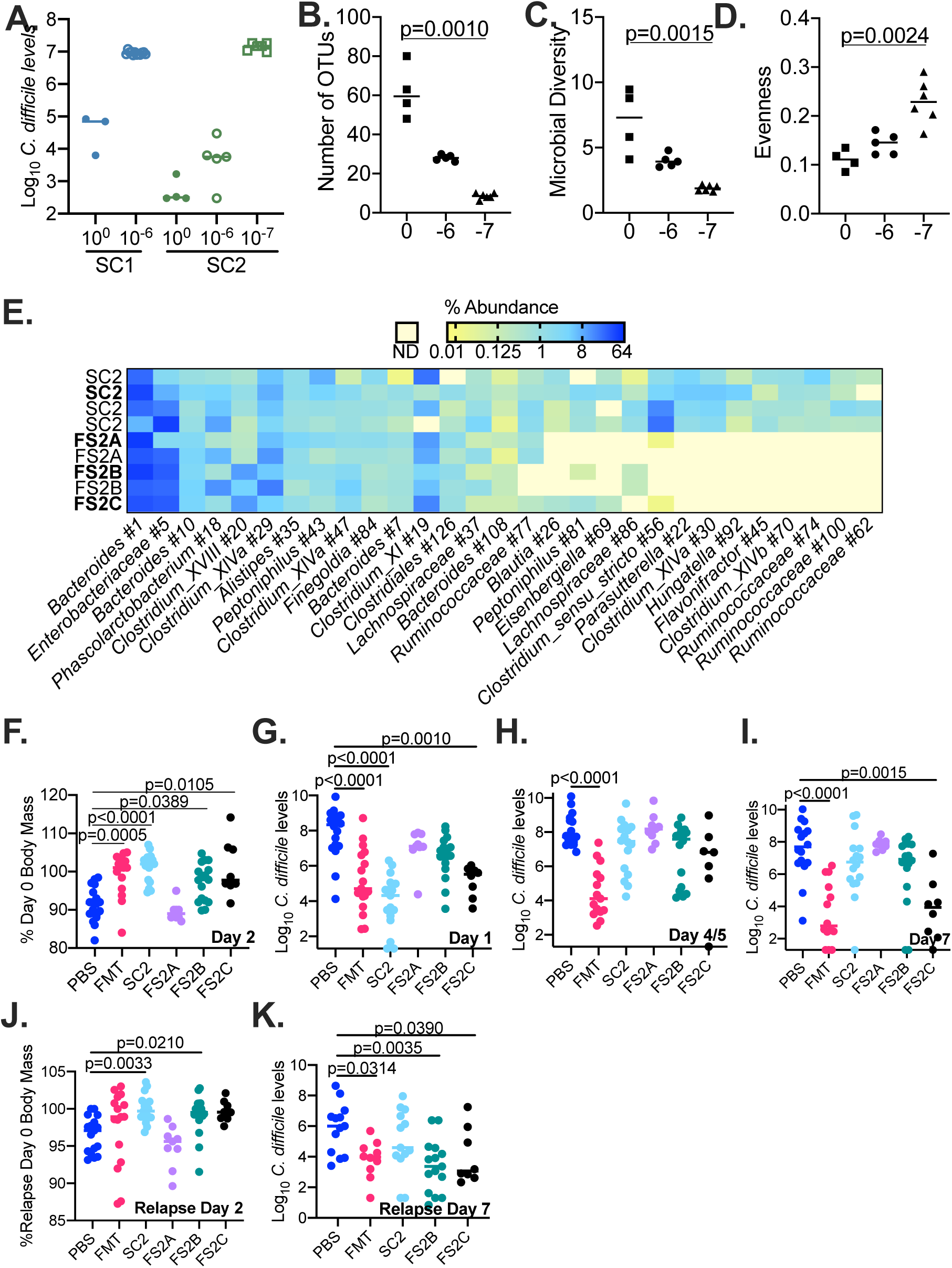
Identification of further simplified microbial communities that suppress *C. difficile* in MBRA and ^HMb^mouse models of CDI. (A) Plot of log_10_ *C. difficile* levels on final day in culture with re-cultured SC1 (closed blue circles) and SC2 (closed green circles) and in re-cultured SC1 and SC2 that were diluted 10^−6^-fold (open circles) and 10^−7^-fold (open squares). Lines represent medians. (B) Number of OTUs, (C) Microbial Diversity (Inverse Simpson Index), and (D) Species Evenness (Simpson Evenness Index) of re-cultured SC2 and 10^−6^ and 10^−7^ diluted communities. Lines represent medians; any significant differences detected (p<0.05) in distributions of 10^−6^ and 10^−7^ diluted communities compared to SC2 communities as determined by one-way Kruskal-Wallis testing with Dunn’s correction for multiple comparisons are reported. (E) Differences in abundance of OTUs present above 0.1% of total sequences in at least two replicate SC2 or FS2 cultures. Samples are indicated to the left of the plot; data in bold-face type are from the cultures shown in 4A; the first SC2 replicate is from the data reported in Figure 2, and the additional SC2, FS2A, and FS2B replicates were collected from bioreactor cultures used to gavage ^HMb^mice in F-K. OTUs are classified to the lowest taxonomic level that could be confidently assigned (≥80% confidence). Yellow represents <0.01% abundance and blue represents ≥ 64% of total sequences as indicated by shading; pale yellow indicates no detected sequences (ND). In(F)-(K), data was collected from ^HMb^mice treated as indicated below the plots. Treatments were administered as described in Figure 3. (F) Percent of day 0 body mass on day 2 of infection. Log_10_ levels of *C. difficile* in mouse feces on day 1(G), day 4/5 (H) or day 7 (I) following initial *C. difficile* challenge. *C. difficile* levels at the mid-point were collected on day 4 or 5 based upon experiment as described in methods. (H) Percent of relapse day 0 body mass on day 2 following induction of relapse with clindamycin IP injection. (I) Level of *C. difficile* detected in feces on day 7 following induction of relapse. Lines represent medians; significant (p<0.05) differences detected in distributions of community-treated mice compared to PBS-treated mice as determined by one-way Kruskal-Wallis testing with Dunn’s correction for multiple comparisons are reported. Longitudinal data from treatments shown in F-K are reported in Fig. S3.

We analyzed the effects of further simplification on community composition and found that the number of OTUs declined from a median of 60 in SC2 cultures to medians of 28 and 9 in 10^−6^ and 10^−7^ diluted communities (Fig. 4B). In addition, microbial diversity decreased (Fig. 4C) and species evenness increased (Fig. 4D) with increasing levels of dilution. The majority of OTUs lost through dilution were classified in the order *Clostridiales* (Fig. 4E). We selected three further simplified communities (FS; FS2A, FS2B, and FS2C) to test for their ability to inhibit CDAD in ^HMb^mice.

### FS2B and FS2C suppress CDAD in ^Hmb^mice

We used the same experimental approach outlined in Fig. 3 to test FS2A, FS2B, and FS2C. As controls, we administered ^HMb^mouse FMT, SC2, and PBS. Similar to our initial study, PBS-treated mice exhibited body mass loss following challenge with *C. difficile* and this was prevented by treatment with ^HMb^mouse FMT (Fig. 4F). Treatment with SC2 also prevented body mass loss, which contrasted with prior results in which SC2-treatment was partially protective. Changes in efficacy could be due to the shifts in microbial composition upon re-culturing of SC2 (Fig. 4E). FS2B and FS2C-treated mice were also protected from CDAD, whereas mice treated with community FS2A lost body mass at a level similar to PBS-treated mice (Fig. 4F).

PBS-treated mice shed *C. difficile* in feces at similarly high levels across all time points tested (Fig 4G-I). In SC2 and FS2C-treated mice, *C. difficile* levels were significantly lower than PBS-treated mice on day 1 of infection, rose by day 4/5 of infection than in SC2 and declined on day 7 of infection. FS2A and FS2B-treated mice showed little reduction in levels of *C. difficile-*shedding. FMT-treated mice had significantly lower levels of *C. difficile* than PBS-treated mice at all time points.

We also tested inhibition of recurrent CDI. Following induction of relapse, we observed a ∼3% median reduction in body mass in PBS-treated mice on day 2 following IP (Fig 4J). Mice treated with SC2, FS2B, or FS2C showed <0.5% median body mass loss, whereas FS2A-treated mice exhibited ∼5% median body mass loss. ^HMb^mouse FMT-treated mice exhibited ∼1% decrease in body mass. We also observed more rapid clearance of *C. difficile* in FS2B and FS2C-treated mice (Fig. 4K).

### Treatment with simplified communities has persistent effects on the fecal microbiome

We analyzed communities from mouse fecal samples on days 1, 4 or 5, 7 and relapse days 0, 2, and 7. Sequence data obtained from mice were pooled with bioreactor data to facilitate tracking of bacteria present in simplified communities in treated mice. We found that sequences clustered at ≥99% ANI (Table S1) provided greater resolution of OTUs distinct to *in vitro*-cultured simplified communities and FMT-treated ^HMb^mice than sequences clustered at ≥97% ANI (Table S2). With sequences clustered at ≥99% ANI, 90% of OTUs found in FMT-treated ^HMb^mice were not detected in *in vitro* cultures and likely represent endogenous bacteria. Similarly, 81% of OTUs found in *in vitro* cultures were not detected in FMT-treated ^HMb^mice. Subsequent analyses used OTUs clustered at ≥ 99% ANI.

The number of OTUs detected on day 1 following infection was low across all treatment groups (Fig. 5A); OTU levels were 25-50% lower than those observed in the FMT sample collected from mice not treated with antibiotics (Fig. 5A, ^HMb^mouse). Treatment with FMT partially restored OTU abundance and microbial diversity on Day 1; full recovery to levels observed in untreated mice was not observed until Day 4/5 during infection. In PBS-treated mice, the median number of OTUs detected in fecal samples increased over time but did not return to the levels detected in untreated mice. Treatment with SC2, FS2C, or FS2B significantly increased the number of OTUs detected on Day 1 compared to PBS-treated mice. Later increases in OTU abundance in FS2C-treated mice paralleled FMT-treated mice treated. For SC2 and FS2B-treated mice, OTU abundance increased over time but not to the extent observed in FMT-treated mice. Neither OTU abundance nor microbial diversity were significantly different between FS2A-treated and PBS-treated mice over the first week of infection. Treatment with FMT, SC2, FS2B and FS2C also significantly increased microbial diversity compared to PBS-treated mice (Fig. 5B).

**Fig 5.**
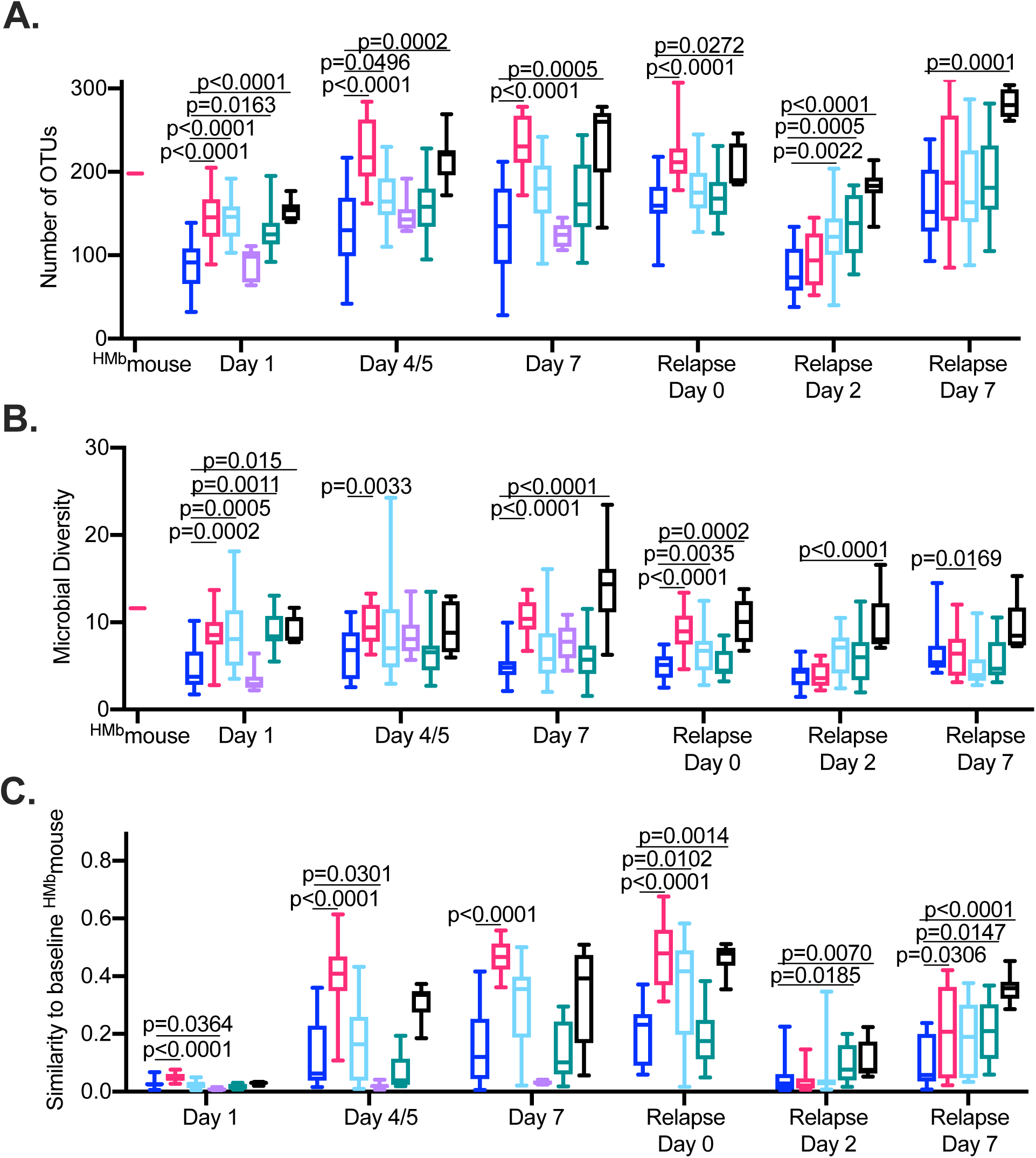
Treatment with SC2 and FS2C restores microbial diversity and shifts microbiome composition towards baseline state observed in ^HMb^mice not treated with antibiotics. 16S rRNA gene sequence data was obtained from bacteria present in the feces of mice treated with FMT (magenta), SC2 (light blue), FS2A (violet), FS2B (teal), FS2C (black) or PBS (dark blue) on days 1, 4 or 5, and 7 following initial *C. difficile* infection and day 0, 2, and 7 relative to initiation of relapse with Clindamcyin IP. (For FS2A-treated mice, 16S rRNA gene sequence data was obtained only from samples collected during initial infection for FS2A-treated mice). 16S rRNA sequence data was also obtained from a pooled fecal sample collected from ^HMb^mouse not treated with antibiotics that was used for FMT administration. (A) Number of OTUs and (B) Microbial diversity (Inverse Simpson Index) measured in sample collected from ^HMb^mice not treated with antibiotics used for FMT administration (^HMb^mouse) and in samples collected from treated mice at time points indicated below graph. (C) Similarity to baseline ^HMb^mouse sample used for FMT administration measured in samples at time points indicated below graph. Boxes represents the interquartile ranges, horizontal lines indicate the medians, and vertical lines indicate the ranges of data collected from replicate mouse samples at each time point. Significance of differences in microbe-treated compared to PBS-treated animals at each time point were evaluated with one-way Kruskal-Wallis testing with Dunn’s correction for multiple comparisons; p-values <0.05 are reported.

We also calculated the similarities in community composition between the baseline ^HMb^mouse sample not treated with antibiotics and communities in the feces of treated mice (Fig. 5C). In FMT-treated mice, fecal communities had low similarities to the baseline ^HMb^mouse sample on day 1, but similarities increased by day 7 (Fig. 5C). In contrast, similarities between PBS-treated and the baseline untreated ^HMb^mouse sample were significantly lower than FMT-treated mice through relapse day 0. Compared to PBS-treated mice, FS2C and SC2-treated mice exhibited an accelerated return towards the baseline microbiome composition. FS2B-treated mice exhibited a return to baseline microbiome that paralleled PBS-treated mice. In contrast, FS2A-treated mice exhibited significantly reduced recovery of microbiome composition compared to PBS-treated mice, indicating FS2A treatment may suppress recovery of the fecal microbiome.

### Treatments shift composition of endogenous bacteria

We identified 98 OTUs that were significantly enriched or depleted in treatments that accelerated microbiome recovery (FMT, FS2C, SC2) compared to treatments with more prolonged disruption (PBS, FS2A, FS2B) for at least one of the time points tested (Table S3). We focused on OTUs with the largest predicted effect sizes (LDA ≥ 3; Fig. 6). Three OTUs, *Erysipleotrichaceae* #4, *Bifidobacterium* #12, and *Bacteroidales* #19 were significantly enriched in the mice FS2C, SC2 and FMT-treated mice at all time points. *Erysipelotrichaceae* #8, *Blautia* #31 and *Clostridium* XIVa #80 were enriched in FS2C, SC2, and FMT-treated mice on day 1, whereas several *Porphyromonadaceae* OTUs (#6,#7,#10,#75, #39, #48) as well as three Firmicutes OTUs (*Erysipelotrichaceae* #17, *Lachnospiraceae* #109, and *Olsenella* #48) were enriched in FS2C, SC2, and FMT-treated mice at later time points. On day 1 of infection, *Peptostreptococcaceae* #35 and *Enterococcus* #28 were enriched in the feces of PBS, FS2A, and FS2B-treated mice on day 1 of infection, whereas *Bacteroides* #3, *Bacteroides* #9, and *Parabacteroides* #1 were enriched during the later stages of infection. These results demonstrate that return towards the baseline microbiome configuration correlates with restoration of members of multiple phyla (*Bacteroidetes, Firmicutes* and *Actinobacteria*) whereas continued disruption correlates increased abundance of *Bacteroides* OTUs and a *Peptostreptococcaceae* OTU that is likely *C. difficile*.

**Fig 6.**
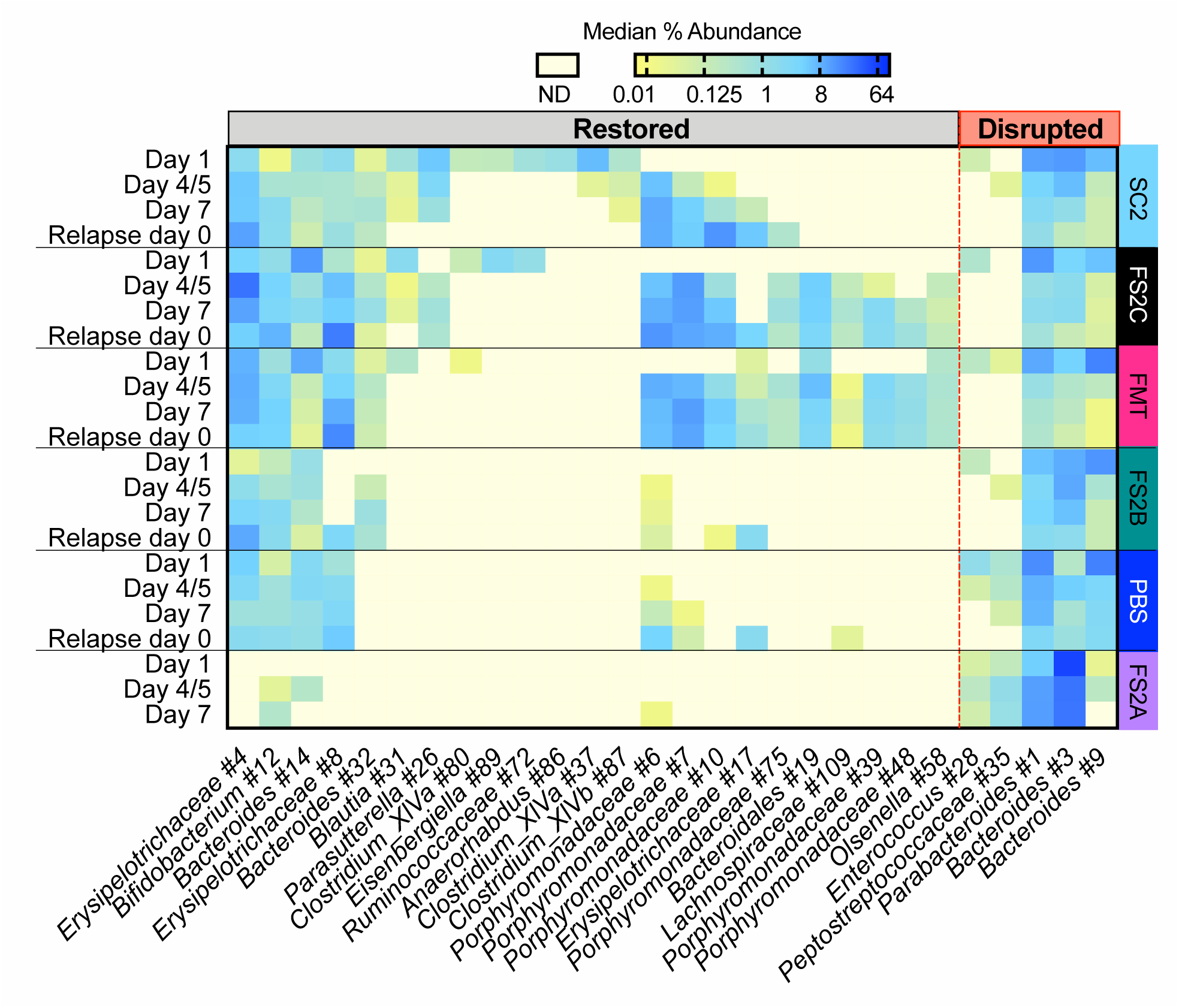
Treatment with simplified communities alters recovery of endogenous microbes. We used LEfSe to identify significantly enriched or depleted taxa between treatments that accelerated microbiome recovery in Fig. 5C (FMT, FS2C, SC2; restorative) and treatments with more prolonged disruption (PBS, FS2A, FS2B; disruptive). Independent analyses were performed for samples on days 1, 4/5, 7, and relapse day 0; OTUs with LDA values determined by LEfSe ≥ 3 for at least one time point are shown. Intensity of shading correlates to the median percent abundance measured for the treated mice at the indicated time points, with median abundances ≥ 64% shaded dark blue and median values equal to 0.01% shaded dark yellow. Samples in which sequences were below the detection limit (ND, not detected) are shaded transparent yellow. OTUs are classified to the lowest taxonomic level that could be confidently assigned (≥80% confidence). *Peptostreptococcaceae* #35 is likely *C. difficile* as the representative sequence has 100% identity to *C. difficile* and abundance over course of infection correlates well with *C. difficile* levels reported in Fig. 3 and 4. The complete set of OTUs identified by LEfSe are provided in Table S3.

### Bacteria from simplified communities persist in the feces of treated mice

We tracked the fate of OTUs present in *in vitro*-cultured simplified communities over time in mice treated with simplified communities. On day 1 following infection, ∼60% of OTUs present in *in vitro-*cultured simplified communities could be detected in the feces of treated mice (Fig 7A-B). Levels of simplified community OTUs decreased over time, with the lowest percentage (∼21%) detected on relapse day 0. Following induction of relapse, the number of OTUs detected from the original *in vitro*-cultured simplified communities increased to ∼53%. These results indicate that these OTUs had likely persisted below the level of detection and re-emerged when other OTUs declined following clindamycin treatment. OTUs that persisted over time were phylogenetically diverse (Fig. 7C). High levels of a *Phascolarctobacterium* OTU (#22) and three *Bacteroides* OTUs (#3, #11, #16) were detected across all community-treated mice indicating that these bacteria likely engrafted well.

**Fig 7.**
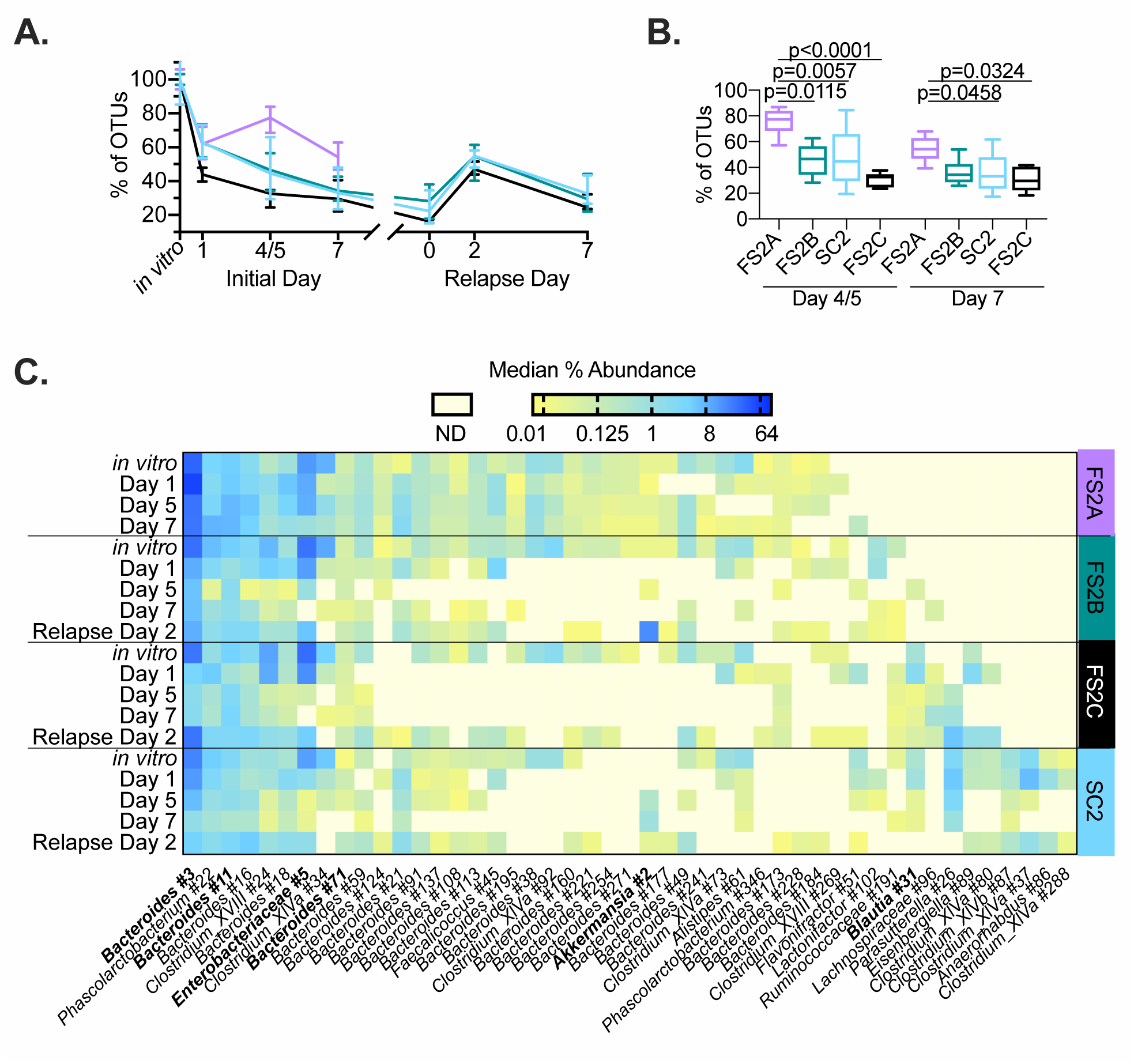
A subset of bacteria from simplified communities persist over time in treated mice. (A) Percent of OTUs in SC2 (light blue), FS2A (violet), FS2B (teal), or FS2C (black) *in vitro* cultures that persist over time in the feces of treated mice. The percent of OTUs detected from each mouse on indicated day relative to median number of OTUs detected in the *in vitro* culture were calculated. Lines represent median values at each time point and error bars represent interquartile ranges. Significance of differences observed between treatment groups at each time point during the initial infection (day 1, 4/5, 7) were evaluated with one-way Kruskal-Wallis testing with Dunn’s correction for multiple comparisons; p-values <0.05 are reported in (B). (B) Data from days 4/5 and 7 re-plotted from (A). Box represents the interquartile range, horizontal line indicates the median, and vertical lines indicate the range of data collected from fecal communities. (C) Percent abundance of OTUs that persist over time in mice treated with SC2, FS2A, FS2B, and/or FS2C. OTUs detected in *in vitro* samples were designated as persistent if the median level in a treatment group on day 7 and/or relapse day 2 sample was ≥0.01%. Intensity of shading correlates to the median percent abundance measured for the treated mice at the indicated time points. OTU labels in bold-face type have also been detected in ^HMb^mice as described in Table S1. Abundance data from persistent OTUs at later times during infection (relapse days 0 and 7) and from OTUs abundant in day 0 samples that did not persist over time in treated-mice are presented in Table S4.

The trend for preservation of OTUs from simplified communities followed a different trajectory in FS2A-treated mice. The percent of FS2A OTUs detected increased to 77% on day 4/5 and returned to 54% on day 7 (Fig. 7A); values were significantly higher than those observed in mice treated with other simplified communities (Fig. 7B). Consistent with this observation, several *Bacteroides* OTUs present in *in vitro* cultures of all four simplified communities were only detected in the feces of FS2A-treated mice on day 7. This increased persistence of OTUs from FS2A was also consistent with the delayed return to baseline microbiome composition observed in these mice (Fig. 5C).

### Bacteria originating from simplified communities re-emerge during relapse

We investigated the microbiome changes were associated with relapse and determined that ∼70% of OTUs enriched in the feces of SC2, FS2C and FS2B-treated mice on relapse day 2 likely originated from the *in vitro*-cultured simplified communities (Fig. 8). Approximately 20% of OTUs enriched on relapse day 2 were enriched on day 1 of infection, indicating that the response to relapse was not identical to the initial disruption but shared some similarities.

**Fig 8.**
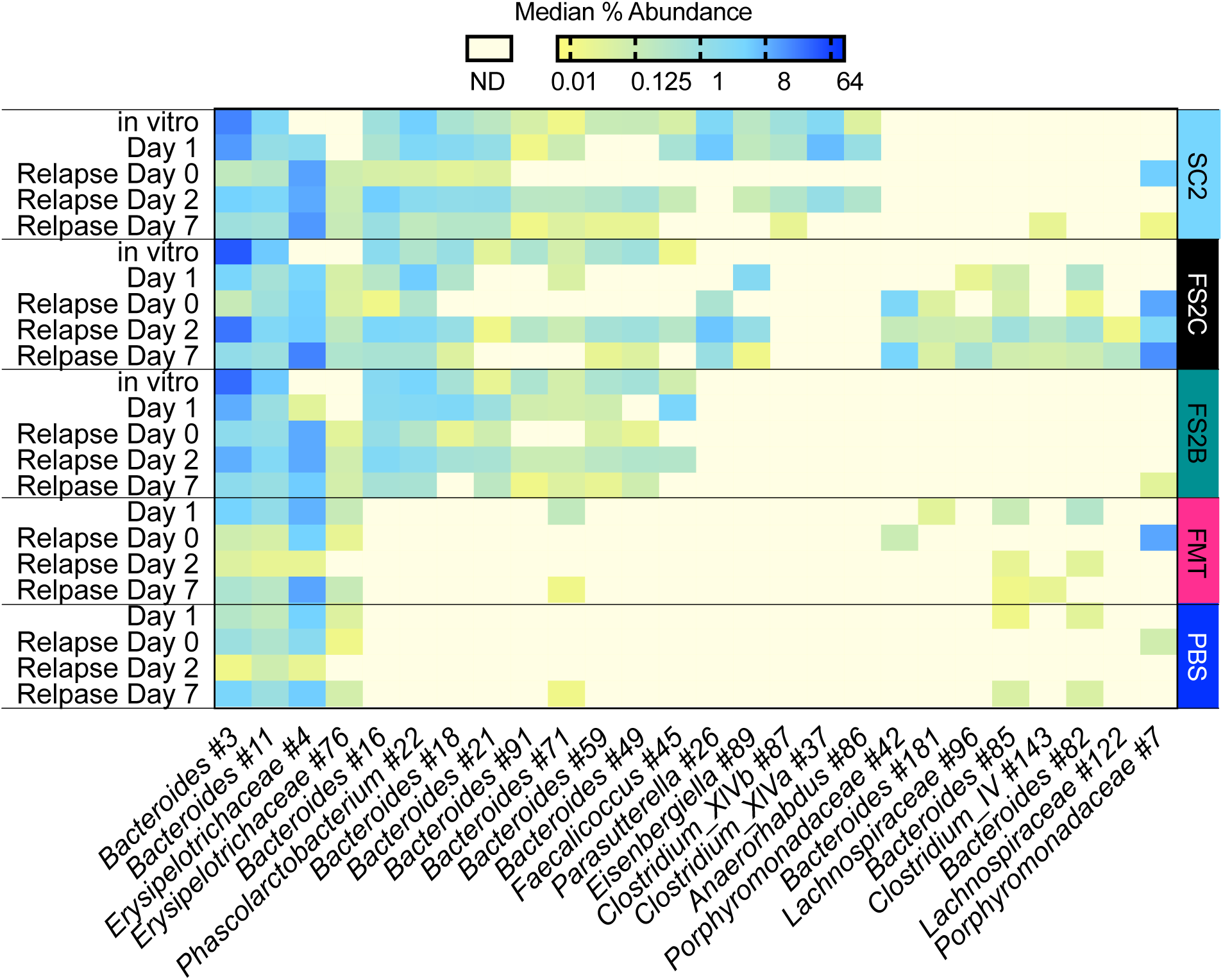
Treatment with SC2, FS2C and FS2B led to distinct microbiome responses during disease relapse. LEfSe was used to identify OTUs that differed significantly between treatment groups in the feces of mice on days 2 and 7 following induction of relapse with clindamycin IP. Independent analyses were performed for samples on relapse day 2 and 7; OTUs with LDA values determined by LEfSe ≥ 3 for at least one time point are shown. The complete set of LEfSe data is shown in Table S5. Median abundance of OTUs in *in vitro* SC2, FS2C, FS2B cultures as well as all the feces of treated mice on day 1 and relapse day 0 samples are included for comparison as described in the text.

## Discussion

We described a new pipeline for identifying and rigorously testing simplified communities with the ability to provide protection from *C. difficile* infection (summarized in Fig. S4). We identified 24 new simplified communities with the ability to inhibit *C. difficile in vitro*. Several of the OTUs detected in these simplified communities were classified into family (*Lachnospiraceae, Ruminococcaceae, Clostridiaceae, Bacteroidaceae*) and genera (*Bacteroides, Clostridium XIVa, Anaerostipes, Coprococcus, Dorea, Roseburia, Blautia*) found depleted in the fecal microbiomes of people who are susceptible to *C. difficile* and restored following FMT treatment (42-46). In contrast, some OTUs were classified into families less often linked to resistance to *C. difficile* colonization (e.g., *Veionella, Eggerthella, Clostridium* XVIII, *Acidaminococcus*) or more correlated with susceptibility to *C. difficile* infection (*Enterococcus, Streptococcus, Escherichia/Shigella*). Thus, the approach we described leads to communities distinct from those based upon predictive ecological modeling and may provide additional insights into *C. difficile* colonization resistance.

By testing simplified communities in a ^HMb^mouse model, we determined that only a subset of the tested communities conferred protection *in vivo.* Treatment with SC1, SC2, FS2B, and FS2C significantly reduced the initial body mass loss associated with severe disease and decreased *C. difficile* loads early in infection, similar to treatment with ^HMb^mouse FMT. While the magnitude of effects varied, we observed a significant negative correlation between *C. difficile* levels on day 1 of infection and body mass on day 2 of infection (Fig. S5), with lower levels of *C. difficile* on day 1 predictive of reduced body mass loss on day 2 of infection. Thus, a potential mechanism for simplified communities to limit the severity of CDI *in vivo* is by delaying the germination or outgrowth of *C. difficile* spores. Similar reductions in *C. difficile* levels on day 1 following infection coupled to ∼50% reduction in body mass loss were reported by Buffie et al (47) when mice were treated with a consortia of four strains. In this case, early reductions in *C. difficile* levels were linked with restoration of secondary bile acid production by *Clostridium scindens*, as well as unknown functions contributed by other members of the simple community. Delaying germination or outgrowth could prevent severe disease by altering the dynamics of the host immune response between pro-inflammatory responses known to cause disease and anti-inflammatory responses that provide protection from *C. difficile* epithelial damage (21, 48, 49).

Comparison of microbiome changes in the feces of mice treated with communities that limit (FS2C, SC2, ^HMb^mouse FMT), partially limit (FS2B) or fail to limit (FS2A, PBS) CDAD may explain some of the observed differences in disease progression. Treatment with FS2C, SC2 or ^HMb^mouse FMT significantly limited body mass loss and altered the levels of *C. difficile* shedding; these communities also exhibited a more rapid return towards the baseline microbiome configuration observed in ^HMb^mice not treated with antibiotics. Return towards baseline was associated with increased abundance of members of the endogenous microbiome, including several *Porphyromonadaceae* OTUs. *Porphyromonadaceae* were found to be depleted in the feces of humans and mice susceptible to *C. difficile* (50, 51). Enhanced restoration of endogenous microbes observed in FS2C and SC2-treated mice could be due to restoration of syntrophic interactions between endogenous microbes and those found in simplified communities and/or suppression of factors (*C. difficile* metabolism (52), innate immune activation (53)) that promote microbiome disruption.

Mice treated with FS2B, a treatment that limited body mass loss during initial infection but did not significantly alter *C. difficile* shedding in feces, exhibited a slower return to baseline microbiome conditions, suggesting that a return to baseline microbiome conditions could be important for *C. difficile* clearance but may not be required to mitigate initial disease severity. Treatment with FS2A, the simplified community treatment that failed to provide protection *in vivo*, was associated with significantly lower levels of restoration of endogenous microbes. Previous reports have indicated that specific probiotic formulations can delay the return to a non-disrupted microbiome configuration due to suppression of endogenous microbes (54); this could also be true for FS2A-treated mice. Further studies are needed to evaluate these hypotheses.

We also found that a subset of OTUs that originated from simplified communities persisted over time in the feces of treated mice. While abundance of these OTUs diminished over time, continued colonization was demonstrated following induction of relapse. Persistence of these simplified community OTUs likely played a key role in limiting susceptibility to recurrent disease. One OTU of note was *Phascolarctobacterium*. A recent study demonstrated that administration of *Phascolarctobacterium* species to cefoperazone-treated mice reduces mortality, possibly by competing with *C. difficile* for succinate in the disrupted GI tract (55). Other OTUs of note include those classified as *Blautia, Ruminococcaceae*, and *Eisenbergiella*. Colonization with members of the *Ruminococcaceae* family and *Eisenbergiella* and *Blautia* genera was correlated with a 60% reduced risk for CDI in allogenic hematopoieitic stem cell patients (56). Our results are also consistent with a previous study of microbiome restoration following FMT in human patients that found a balance between engraftment of donor bacteria, persistence of bacteria present in the feces of infected patients, and emergence of previously undetected bacteria ((57).

Dilution-extinction provided a rapid way to screen communities for the ability to prevent *C. difficile* infection. Development of diverse treatment consortia for CDI is important as *C. difficile* is known to fill different nutritional niches (58) and fecal transplant studies indicate differential engraftment of species between patients treated with the same fecal sample (59). However, further refinement is needed before communities progress to clinical testing. Isolation of individual strains from simplified communities prior to community reassembly and efficacy testing will ensure the identity of the treatment consortia. In spite of these limitations, the approaches outlined in this study represent a significant advance in the throughput of testing for simplified communities to limit *C. difficile* infection and could potentially be adapted to identify simplified communities to treat other diseases linked to microbiome disruption.

## Methods

### Fecal samples, bacterial strains, and cultivation conditions

Fecal samples were provided by anonymous subjects between the ages of 25-64 who self-identified as healthy and had not consumed antibiotics for at least 2 months or probiotics for at least 2 days prior to donation. Fecal samples were prepared as described (60). The previously described ribotype 027 isolate *C. difficile* 2015 was used for all experiments (60). All cultivation was performed at 37°C under an atmosphere of 5% H_2_, 5% CO_2_, and 90% N_2_.

### Identification of simplified communities through dilution/extinction

Fecal samples were prepared and inoculated into MBRAs containing bioreactor media 3 (BRM3) (61) as described (40). Fecal communities equilibrated for 16 hr in batch growth containing before initiation of continuous flow at a flow rate of 1.875 ml/hr (8 hr retention time). After 5-6 days of flow, an aliquot was removed for determination of cell concentration through serial dilution and plating on BRM3 agar. After 8 days, a sample was removed from each reactor, diluted to final concentrations of ∼3 × 10^4^ cells/ml (10^−4^ dilution) or 3 × 10^3^ cells/ml (10^−5^ dilution) in BRM3. 1 ml of each dilution was used to inoculate 5-6 sterile bioreactors containing 15 ml of sterile BRM3/dilution. After 3 days under continuous flow, aliquots were removed from diluted communities for sequencing and for cryopreservation with 15% glycerol or 7.5% DMSO. One day later, communities were challenged with 10^4^ *C. difficile* cells as described (60); *C. difficile* levels in reactors were determined through selective plating on TCCFA agar with 20 μg/ml erythromycin and 50 μg/ml rifampicin as described (60). For repeat cultivation from cyropreserved stocks, stocks were thawed and 300 μl were used to inoculate triplicate reactor vessels containing 15 ml of sterile BRM3. Communities were grown in batch for 16 hrs, then with continuous flow for four days prior to challenge with *C. difficile* as described above.

### Further simplification of simplified communities 1 (SC1) and 2 (SC2)

1 ml stocks were thawed and used to inoculate an empty reactor vessel. Flow of sterile BRM3 was initiated and allowed to fill the reactor at a flow rate of 1.825 ml/hr. After continuous flow cultivation for three days, cell concentrations were determined as described above. Two days later, aliquots of cells were removed and diluted in sterile BRM3 to a final concentration of 250 cells/ml (10^−6^ dilution) or 25 cells/ml (10^−7^ dilution). 1 ml aliquots of cells were used to inoculate 5 (10^−6^ dilution) or 6 (10^−7^ dilution) empty, sterile reactors, which were allowed to fill with sterile media as described above. After two (SC2) or three (SC1) days of flow, aliquots were removed for sequence analysis and cryopreservation. One (SC1) or 15 (SC2) days later simplified communities were challenged with 10^4^ vegetatively growing *C. difficile* cells and levels of *C. difficile* persisting in reactors over time were determined through selective plating.

### Cultivation of simple communities for treatment of ^Hmb^mice

65 ml bioreactors were prepared as previously described (62). Sterile, empty bioreactors were inoculated with 1 ml of thawed stocks and allowed to fill with sterile BRM3 medium at a flow rate of 8.125 ml/hr. Communities were cultured with flow for 6-8 days before 10 ml aliquots of culture were removed, centrifuged at 800 X g for 10 min and resuspended in 1 ml anaerobic phosphate buffered saline for delivery to mice. Cell densities of reactor communities were determined through selective plating on BRM3 agar; mice received doses of cells ranging from 5 × 10^8^ – 2 × 10^9^ cells freshly prepared from reactors on three subsequent days.

### Preparation of ^HMb^mouse FMT and human FMT material

Fecal samples were collected from 6-10 week-old male and female mice, pooled and resuspended in anaerobic PBS at 20% w/v. Samples were vortexed for 5 min, then centrifuged at 200 X g for 2 min. Each mice was treated with 100 μl of fecal slurry. Our human FMT preparation was prepared as described (63).

### Treatment of ^HMb^mice with PBS, human FMT, ^HMb^mouse FMT or simplified communities

As outlined in Fig. 3A, antibiotics (60) were administered in the drinking water to 6-10 week-old male and female mice. Mice were treated with 100 μl of PBS, ^HMb^mouse FMT, human FMT or cells from simplified communities via orogastric gavage on three subsequent days. Clindamycin (10 mg/kg) was administered via intraperitoneal injection. Mice were challenged with 5 × 10^4^ spores of *C. difficile* 2015. Three mouse experiments were performed. Mice in experiment 1 were treated with PBS, ^HMb^mouse FMT, human FMT, or SC1-SC4 (n=9 mice/treatment group except SC4 (n=8)). Mice in experiment 2 were treated with PBS, ^HMb^mouse FMT or SC2, FS2A, or FS2B (n=9 mice/treatment group except PBS (n=10)). Mice in experiment 3 were treated with PBS, ^HMb^mouse FMT or SC2, FS2B, or FS2C (n=8 mice/treatment group). In experiments 1 and 2, ∼100 μl of inoculum from the 3rd treatment was saved for sequencing. Mouse body mass was collected daily from days 0-5 following *C. difficile* challenge then periodically following resolution of severe disease as indicated in figures. Mice that lost greater than 20% body mass from day 0 or showed signs of severe disease as previously described (41) were euthanized. Mouse body mass was also collected on day -2 and day -1 in experiment 1. Relapse was induced 24 (experiment 3), 28 (experiment 1) or 33 (experiment 2) days following initial *C. difficile* infection through IP administration of clindamycin (10 mg/kg). *C. difficile* levels in fecal samples were determined through selective plating (experiment 1) or qPCR (experiments 2 and 3) as described (60) at the time points indicated in the text.

### Analysis of microbial communities through 16S rRNA gene sequencing

Nucleic acids were extracted from mouse fecal samples and inoculum samples using the DNeasy Powersoil HTP Kit (QIAGEN) and from the further simplified SC2 samples using the Powermag Microbiome kit (MoBio). The V4 region of the 16S rRNA gene was amplified from purified DNA or directly from lysed bioreactor samples in triplicate using dual or single indexed primers F515/R806 as described (40, 64). Samples were cleaned quantified and pooled in equimolar concentrations prior to sequencing using the Illumina MiSeq v2 2 × 250 kit as described (40).

All sequence analysis was performed using mothur version 1.35.1. Raw sequencing reads were quality-filtered, aligned to the V4 region of Silva 16S rRNA reference release 132, pre-clustered into sequence groups with <1% sequence divergence, filtered to remove chimeras with uchime, and classified with the Bayesian classifier using rdp database version 16 (≥80% confidence threshold) as previously described with the modifications noted above (40, 65). Sequences were then rarefied to remove those with ≤ 10 reads. Pairwise distance matrices where calculated and sequences were clustered into OTUs with ≥97 and ≥99% ANI. OTUs were classified by the majority consensus rdp taxonomy within the OTU. To better determine the potential identity of *Peptostreptococcaceae* OTU #31 (≥97% ANI) and *Peptostreptococcaceae* OTU #35 (≥ 99% ANI), representative sequences from these OTUs were compared to the nr/nt database using BLAST.

Samples were randomly subsampled to 10,000 sequences before determination of alpha and beta diversity measures. Alpha diversity measures (number of observed OTUs, inverse Simpson measure of microbial diversity, Simpson even measure of evenness) were calculated using mothur. Principle Coordinates Analysis of Bray-Curtis dissimilarities between communities were calculated and ordinates were visualized using the Phyloseq package (version 1.30.0 (66)) running in R version 3.61. Statistical significance of clusters were calculated with permutational ANOVA (ADONIS function of vegan version 2.5-6 (40)). Identification of OTUs significantly enriched between treatment groups was determined using the mothur-implementation of LEfSe (40, 65). Mothur was also used to calculate the Bray-Curtis dissimilarities between treated mice and the baseline ^HMb^mouse sample (similarity= 1-Bray-Curtis dissimilarity).

### Ethics statement

Protocols for fecal sample collection were reviewed and approved by the Institutional Review Boards of Michigan State University and Baylor College of Medicine. Animal use was reviewed and approved by the Institutional Animal Care and Use Committee at Baylor College of Medicine (protocol number AN-6675)

### Data visualization and statistical analysis

Unless otherwise noted, data was visualized and statistical analysis was performed using Prism v8.

### Data availability

16S rRNA gene sequence data has been deposited in the sequence read archive ((66)) with accession numbers XXX.

## Acknowledgements

This work was supported by funding from the National Institutes of Allergy and Infectious Disease (AI12152201) to RAB and from the Nebraska Tobacco Settlement Biomedical Research Development Fund to JMA. Analysis was completed utilizing the Holland Computing Center, which receives support from the Nebraska Research Initiative. RAB receives unrestricted research support from BioGaia, AB, consults for Takeda and Probiotech, serves on the scientific advisory board of Tenza, and is a co-founder of Mikrovia.

The authors thank T. Savidge (Texas Children’s Microbiome Center, Baylor College of Medicine) for providing the human FMT sample.

## Supplemental Material

### Figure Legends

**Figure S1. *C. difficile* proliferation in triplicate cultures seeded with initially suppressive communities.** *C. difficile* levels from triplicate reactors (closed circles) re-cultured from cryopreserved simplified communities are plotted with the donor sample designation indicated below the graph and shading as indicated in Figure 2. Open circles indicate levels of *C. difficile* detected in the initial culture reported in Fig 1B. Lines represent medians of all four data points. Dotted lines indicate levels of *C. difficile* that are ≥100 and >10,000 times lower than the maximum *C. difficile* levels reported in Figure 1.

**Fig S2. Simplified microbial communities SC1 and SC2 suppress *C. difficile* in** ^**HMb**^**mouse model of CDI.** Longitudinal data collected from ^HMb^mice that were administered treatments described in Figure 3. (A-D) % of day 0 body mass and (E-I) Log_10_ *C. difficile* levels in feces of mice over the first seven days following infection. (L-O) % of relapse day 0 body mass and (P-S) log_10_ *C. difficile* levels in feces of mice over time following initiation of relapse by IP injection of clindamycin. Data presented are from mice treated with human FMT (A,F,KO), ^HMb^mouse FMT (E,J,N), SC1 (B,G,L,P), SC2 (C,H,M,Q) and SC3 (D,I) Lines indicate medians and error bars indicate interquartile ranges. Values for PBS-treated mice are in dark blue. Treatments that differ significantly from PBS-treated mice were evaluated with one-way Kruskal-Wallis testing with Dunn’s correction for multiple comparisons with p<0.05 indicated by asterisks. Response during relapse was not tested in SC3-treated mice.

**Fig S3. FS2B and FS2C communities suppress *C. difficile* in** ^**HMb**^**mouse model of CDI.** Longitudinal data collected from ^HMb^mice that were administered treatments described in Figure 4 and in the text. (A-E) Percent of day 0 body mass and (F-J) log_10_ levels of *C. difficile* in feces over time during the first seven days following infection. (K-O) Percent of relapse day 0 body mass and (P-S) levels of *C. difficile* in feces over time during the seven days following initiation of relapse by IP injection of clindamycin. Data presented are from mice treated with ^HMb^mouse FMT (A, F, K, P), SC2 (B, G, L, Q), FS2A (C, H, M), FS2B (D, I, N, R) or FS2C (E, J, O, S). Data from PBS-treated mice (dark blue lines) are repeated in each panel for reference. Lines represent median values and error bars represent interquartile ranges. Data points identified as statistically significant in Figure 4 are indicated by asterisks.

**Fig S4. Summary of process used to identify simplified communities with ability to inhibit *C. difficile in vitro* and limit *C. difficile* associated disease *in vivo*.** As described in the text, simplified communities were initially generated through dilution of complex fecal communities and tested for their ability to inhibit *C. difficile* persistence *in vitro.* A subset of original retained ability to inhibit *C. difficile* when re-grown from frozen stocks. Four communities (SC1, SC2, SC3, and SC4) were identified for subsequent *in vivo* testing. Two of these simplified communities, SC1 and SC2 were tested to see if they could be further simplified *in vitro* and retain ability to inhibit *C. difficile.* SC1 lost ability to inhibit *C. difficile* upon dilution but it was retained by SC2. Three further simplified SC2 communities were identified for testing *in vivo* and designated FS2A, FS2B, and FS2C. SC1, SC2, SC3, SC4, FS2A, FS2B, and FS2C were tested *in vivo* in a humanized microbiota mouse model of *C. difficile* infection. SC1, SC2, FS2B, and FS2C provided protection from *C. difficile* associated disease. SC3 and FS2A failed to provide protection. SC4 was toxic to mice prior to *C. difficile* administration.

**Fig S5. Correlation analysis of levels of *C. difficile, C. difficile* toxin, and body mass change during initial infection.** Correlation analysis of % body mass on day 2 of infection relative to *C. difficile* levels on (A) day 1, (B) day 4/5, or (C) day 7 from all mice tested in experiments 1-3. Linear regression formulas and correlation coefficients for each plot are indicated in the corner of each graph. For regression plots, p<0.0001.

### Tables

**Table S1.** Characterization of OTUs clustered at >99% ANI shared by simplified communities and ^HMb^mice.

**Table S2.** Characterization of OTUs clustered at > 97% ANI shared by simplified communities and ^HMb^mice.

**Table S3.** Abundance of OTUs that differed significantly between treatments that restored microbiome diversity towards baseline (FMT, FS2C, SC2) and those that did not (PBS, FS2A, FS2B).

**Table S4.** Persistence of OTUs present in SC2, FS2A, FS2B, and FS2C over time in treated ^HMb^mice.

**Table S5.** Abundance of OTUs in control and microbe-treated ^HMb^mice that differ significantly between treatments.

